# The multifaceted process of human coronary atherosclerotic cap destabilisation

**DOI:** 10.1101/2024.07.21.604507

**Authors:** L.E. Bruijn, N. Fonseca Neves, C.M. van Rhijn, J.F. Hamming, AJ van den Bogaerdt, J.H.N. Lindeman

**Affiliations:** Dept. of Surgery, Division of Vascular Surgery, Leiden University Medical Center (LUMC); ETB-BISLIFE, Heartvalve Department, Beverwijk, The Netherlands

## Abstract

**Introduction:** Plaque rupture is the primary trigger of the acute clinical manifestations of atherosclerotic disease. So far, factual insight in the processes leading up to cap destabilization is largely missing. In order to overcome this knowledge gap, a pseudo-timeline of atherosclerosis progression was established in order to systematically map the qualitative changes in cap characteristics during lesion progression and destabilization.

**Material and Methods:** A pseudo-timeline was created by randomly selecting preclassified (revised AHA classification, at least 10 per stage) left coronary artery FFPE specimens obtained during tissue donation (aortic valve procurement). Qualitative changes were visualized by (immuno)histochemistry, immunofluorescence and confocal microscopy. Scoring was performed by two observers using semiquantitative scoring estimates.

**Results:** The median age of the donors was 56 years (IQR 51.5-59), and 67% of the patients was male. Movat staining indicated a consistent pattern of cap formation, maturation and destabilization. A distinctive cap emerged in the early fibroatheroma stage of progressive atherosclerosis. Disease progression was accompanied by profound fibrotic changes in the gap, and a progressive presence of inanotic (nutritional deprivation leading to dissolution) mesenchymal cells. Plaque rupture was preceded by thinning of the collagen fibers and accumulation of foam cells in the central portion of the thin cap. No evidence was found for a direct involvement of neovascularization in the destabilization process.

**Conclusion:** The pseudo-time line of atherosclerotic lesion development characterizes the development of an unstable cap as a degenerative and fibrotic process with progressive exhaustion of the mesenchymal cell population. This study provides a rationale for the limited efficacy of medical strategies aimed at plaque stabilization.

## Introduction

Atherosclerosis is a chronic, if not lifelong, process that paradoxically generally manifests as an acute ischemic event.^1^ This paradox is thought to reflect the very nature of the disease. Atherosclerotic lesion formation is a gradual process that is driven by intimal LDL deposition and foam cell formation. Progressive cholesterol accumulation and foam cell death results in formation of the classic atherosclerotic lesion: a cholesterol crystal-rich necrotic core that is shielded from the blood stream by a fibrous cap. Ultimately, destabilization (rupture) of the cap exposes the necrotic core contents to the blood stream, potentially triggering thrombus formation, and an acute vascular occlusion (infarction).^2^

Notwithstanding the pivotal role of cap destabilization as the trigger of acute ischemic events^3^, factual insight in the processes leading up to cap destabilization is limited. As a consequence qualitative measures of cap stability are not included in the prevailing grading systems. For example, the AHA classification^4^ exclusively addresses presence or absence of a “surface defect” as a dichotomous measure of cap status. The modified AHA classification system as proposed by Virmani et al^5^, introduced the vulnerable lesion type (thin cap fibroatheroma) in which cap stability is graded by cap thickness (≥ or< 65 μm) as a dichotomous marker of plaque stability. Qualitative aspects such as the extent of macrophage infiltration^6^, or the extent of smooth muscle cell loss^7^, have been reported in isolation. Yet, as this point, a more refined classification of plaque stability required to systematically study the destabilization process, let alone the full picture of the destabilization process, is missing. This knowledge gap partly relates to the fact that most experimental atherosclerosis models do not progress to the classic vulnerable lesion type.^8^ On the other hand, clinical (human) studies generally rely on surgical or post-mortem specimens, in which the distinction between ‘stable’ and ‘unstable’ lesions is made on basis of the clinical manifestation(s), such as infarction or TIA. Hence, in the context of surgical specimens, the term unstable plaque often refers to a post-event (rupture) situation rather than to the process of plaque destabilization.^8^ As a consequence, current understanding of the cap stabilization process is largely based on theoretical axioma’s based on extrapolation of the available data.

Availability of a large tissue repository of human left coronary arteries from tissue donors (heart valve collection for tissue donation) with atherosclerotic lesions spanning the full spectrum of atherosclerotic disease, allowed us to construct a pseudo-timeline of the cap destabilization process. Based on this timeline, the qualitative changes in cap characteristics during lesion progression and destabilization were systematically mapped. This observational study characterizes plaque rupture as a result of the progressive fibrotic changes in the cap.

## Methods

### Patients and tissue sampling

Use of donor material for scientific purposes is approved by the Medical and Ethical Committee of the Leiden University Medical Center, The Netherlands. Sample collection and handling was performed in accordance with the guidelines of the Medical and Ethical Committee of the Leiden University Medical Center, with the code of conduct of the Dutch Federation of Biomedical Scientific Societies, and in accordance with the principles of the Declaration of Helsinki.

Due to national regulations, only data relevant for transplantation was available for research (http://www.federa.org/codes-conduct). Specific information on risk markers such as cholesterol or CRP levels were not available for the donors. Information on smoking (history) relied on information provided by relatives.

This study includes material from a tissue repository of more than 900 classified (modified AHA classification according to Virmani^5^) proximal left coronary artery (LCA) segments from as many Dutch post-mortem heart valve donors who donated in the period 2011–2021. The current age limit for valve donation is 65 years for men and 70 years for women (65 years until 2016). As such, the maximum age of individuals in the study is 70 and the number of specimens from individuals over 65 years is limited. Further main exclusion criteria for donation include: (a history of) malignancy, sepsis and/or risk of transferable disease (hepatitis, prions etc.), and/or (suspected) connective tissue disorders or vasculitis/myocarditis. Donation criteria further exclude men over 50 years with a history of diabetes, or chronic kidney disease, Chronic Obstructive Pulmonary Disease or Abdominal Aortic Aneurysm.

Donor screening and acceptance was performed by the Dutch Transplant Foundation (Leiden, The Netherlands), and all donors gave permission for research. Valve dissection for homograft valve transplantation was performed within 40 hours post-mortem by ETB-BISLIFE (Heart Valve Department, Beverwijk, The Netherlands) by carefully removing the aortic and pulmonary valve from the intact heart. Both valves were freed from the surrounding pericardial fat and banked for transplantation. The pericardial fat tissue containing a segment of the proximal LCA (approximately 2-3cm) was formaldehyde fixed and used in this study. This collection procedure did not interfere with the actual valve dissection or the pathological analysis of the heart necessary for the final release of the banked valves.

55 tissue blocks with lesions types ranging from Pathological Intimal Thickening (PIT) to Plaque Rupture (PR) were randomly selected from the repository and the material used in this study. Of note, sections representing PR are rare and all available cases (n=14) were included in this study.

### Tissue processing

Formaldehyde-fixed tissue was decalcified (Kristensen’s solution), divided in consecutive 5 mm segments, and each segment was embedded in paraffin. Decalcification does not interfere with (calcium) scoring, as histological footprints of earlier calcium deposits (hydroxyapatite) remain present after the process of decalcification (identified as brown and dark purple deposits in Movat staining^9^). Four μm sections were prepared for each individual segment and Movat pentachrome staining for atherosclerosis grading (revised classification of the American Heart Association (AHA) as proposed by Virmani et al.^5^) was performed for each segment. Movat stainings were complemented with Picro Sirius Red (Abcam; ab150681; one hour, washed in acetic acid) and H&E stainings for respectively their superior reflection of collagen matrix, and nuclear counts and cell morphology.

### Single-labelling immunohistochemical staining

Primary antibodies (summarized in **Table 1**) were diluted in 1% PBS/BSA and incubated overnight at room temperature. Heat-induced antigen retrieval was performed for 10 minutes and endogenous peroxidase activity was blocked with a 20-min incubation of 0,3% hydrogen peroxide. The Envision/DAB (Dako, Glostrup, Denmark) system was used for visualization. Nuclei were counter stained by Mayer’s hematoxylin (Merck Millipore, The Netherlands). Microvessels were visualized using Biotinylated Anti-Ulex Europaeus Agglutinin l (Vector Laboratories Inc, Burlingame, United States), Vectastain ABC HRP Kit (Amsterdam, the Netherlands) and DAB.

**Table 1:**
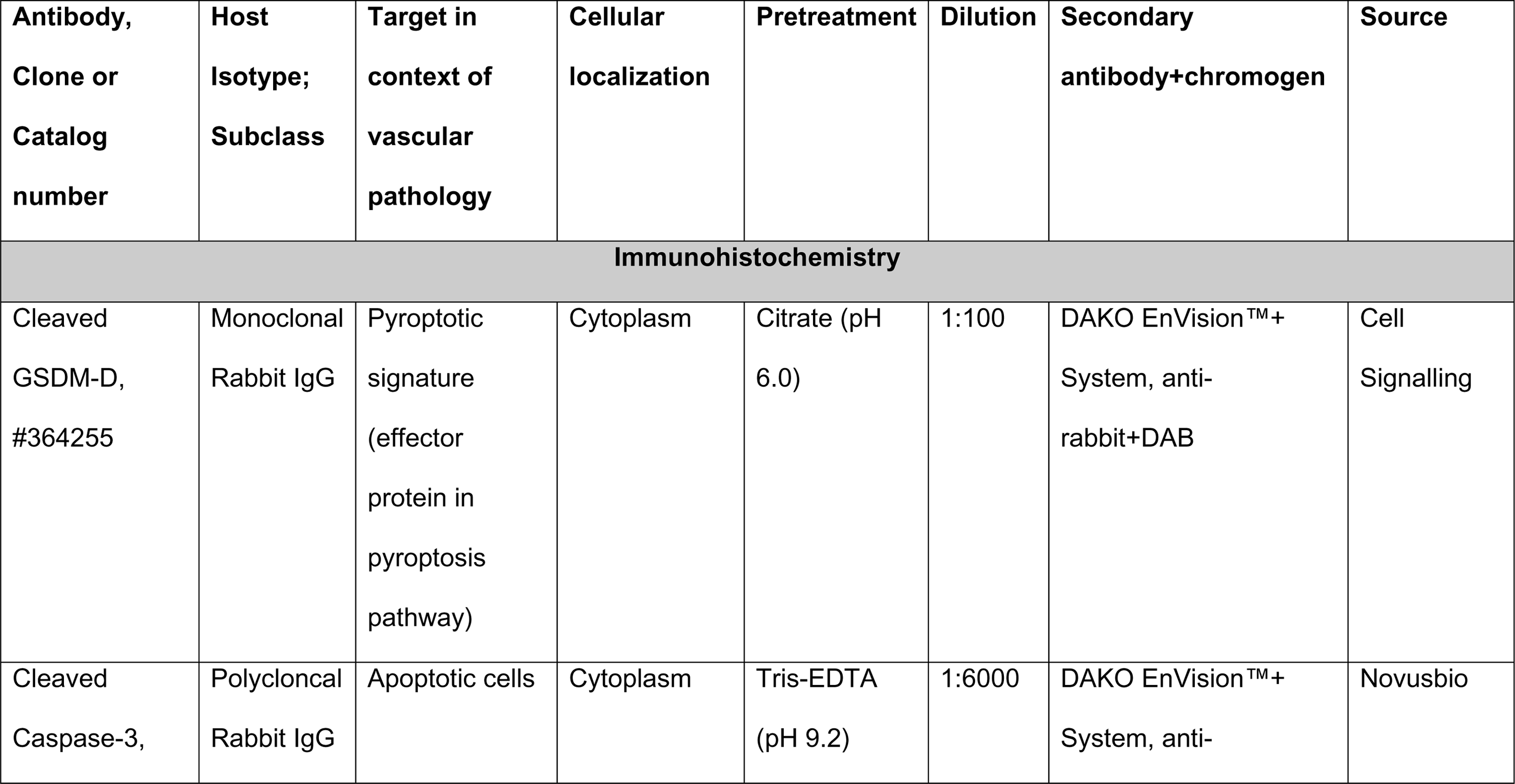
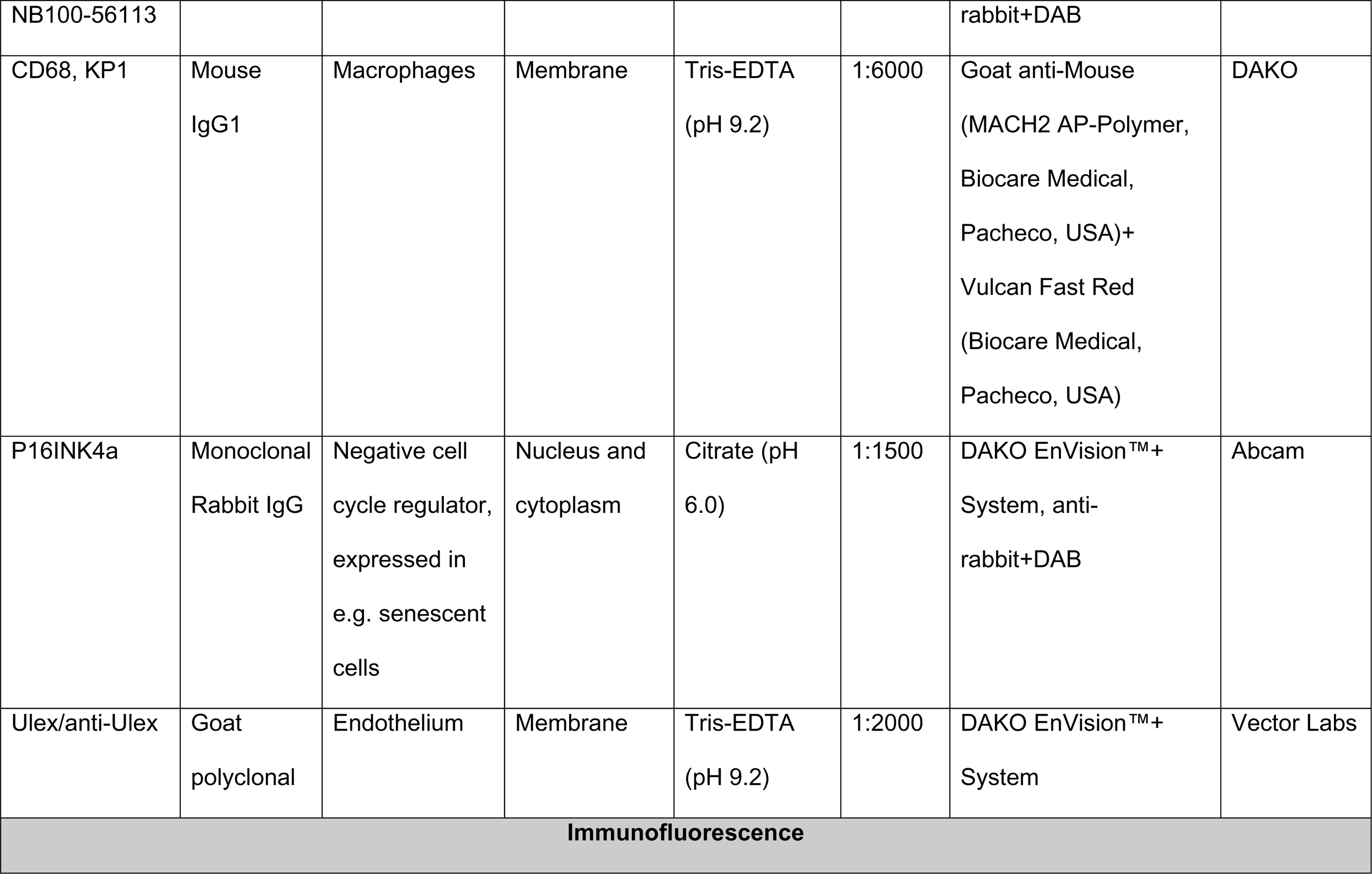
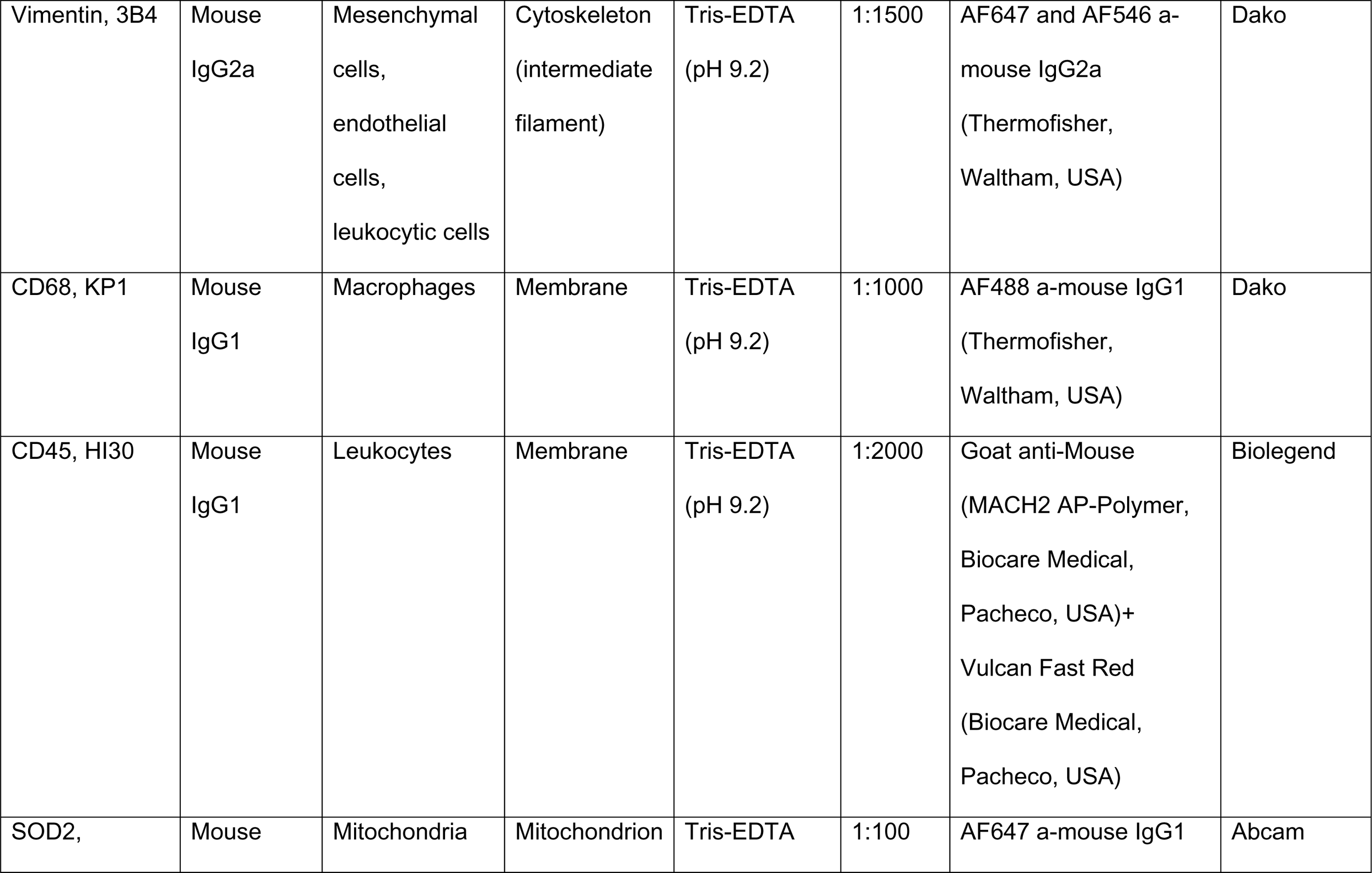
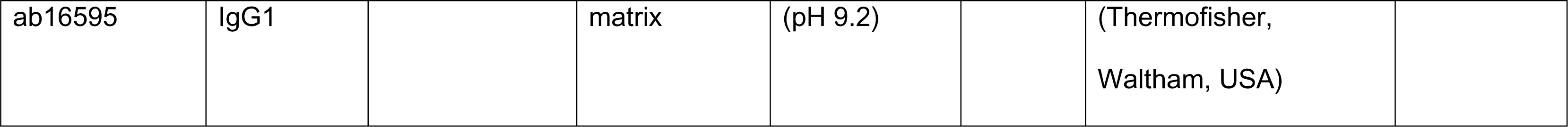
Antibodies used in the present study.

### Double-labelling immunohistochemical staining

Double-labeling IHC staining (**Table 1**) was performed by sequential single labeling IHC. A second heat-induced antigen retrieval after the first chromogen staining was used to inactivate the previous signal. All epitopes resisted the second heat-retrieval. The alkaline phosphatase labelled secondary antibody (MACH2 AP Polymer, Biocare Medical, Pacheco, USA) with subsequently Vulcan red (10 min, dilution 1:50; BioCare Medical, Pacheco, USA) and the Envision/DAB (Dako, Glostrup, Denmark) system were used for visualization. Double stained slides were not counterstained.

### Triple-labeling immunofluorescence staining

Slides were washed with Triton X-100 (Abcam, Cambridge, UK) 0.1% in PBS for 10 minutes prior to incubation of primary antibodies (summarized in **Table 1**) and heat-induced antigen retrieval was performed for 10 minutes. For the CD45/CD68/Vimentin staining, CD45 staining was first performed as a single staining using Vulcan Fast Red for visualization. Next, a second heat retrieval was performed, and the sections incubated overnight with CD68/Vimentin. Corresponding Alexa Fluor secondary antibodies (dilution 1/200; Thermofisher, Waltham, USA) were incubated for 60 minutes at room temperature. Nuclei were visualized using Hoechst 33342 (Thermofisher, Waltham, USA). Slides were mounted using EverBriteTM Mounting Medium (Biotium, Fremont, USA) and stored at 4°C until analysis. Vulcan Red fluorescence was visualized using a Texas Red Filter (542-582 nm).

### Imaging

Immunohistochemistry images were captured by means of a digital microscope (Philips IntelliSite Pathology Solution Ultra-Fast Scanner, Philips Eindhoven, the Netherlands).

Images of triple IF stainings were acquired using the Panoramic MIDI Digital Slide Scanner (3D HISTECH Ltd, Budapest, Hungary) and analyzed with CaseViewer software (3D HISTECH Ltd). Minor linear adjustments (brightness and contrast) were performed. Non-linear adjustments were not performed.

Confocal images were acquired with the Zeiss Confocal Airyscan LSM900 (Zeiss, Oberkochen, Germany). We used 20×/0.8-NA (numerical aperture), 40×/1.4 NA and 63×/1.40-NA oil-immersion objective lenses. The lasers included a 405 nm, 488 nm, 561 nm, and 640 nm. Z-stacks were created in Zen Blue (Zeiss, Oberkochen, Germany). 3D images were acquired by Imaris x64 software (Oxford Instruments, Abingdon, Oxfordshire).

### Definition of the cap and shoulder regions and cap thickness measurements

A schematic representation of the different aspects of the cap is provided in **Supplemental Figure 1**. The fibrous cap was defined as the tissue between the necrotic core and the vessel lumen, restricted by the shoulder regions. The shoulder regions were defined as the tissue between a projected perpendicular line between the outer border of the necrotic core and the directly underlying media down to the vessel lumen; and the border between the lesion and non-lesional (“normal”) intima.

### Statistics

Scoring of marker expression was performed by 2 observers using semiquantitative scoring estimates (ie, 80% positivity). Cap thickness was measured in two areas: in the mid section of the cap, and in the border between the cap and the shoulders. The mean of the readings for the left and right shoulders was used to define cap thickness between the cap and the shoulders.

## Results

### Donors included for pseudo atherosclerosis timeline

A pseudo time-series was created by randomly selecting preclassified (adapted AHA classification according to Virmani^5^) left coronary artery specimens from the tissue repository. The selection represents the successive stages of progressive atherosclerosis:^5^ plaque initiation, maturation and destabilization, viz. Pathological Intimal Thickening, (PIT, n=10), Early Fibroatheroma (EFA, n=10), Late Fibroatheroma (LFA, n=10), Thin Cap Fibroatheroma (TCFA, n=11) and Plaque Rupture (PR, n=14).

Clinical characteristics of the cases included in the timeline are summarized in **Table 2**. The median age was 56 years (IQR 51.5-59), and 67% of the patients was male. The minority of the patients was treated with statins (20%) or antihypertensives (31%). 87% of the patients died of a cardio- or cerebrovascular cause.

**Table 2:** Patient characteristics (n=55)

|  | Pathological Intimal Thickening (PIT),<br>*n=10* | Early fibroatheroma (EFA),<br>*n=10* | Late Fibroatheroma (LFA),<br>*n=10* | Thin cap fibroatheroma (TFCA),<br>*n=11* | Plaque Rupture (PR),<br>*n=14* |
| --- | --- | --- | --- | --- | --- |
| Age, median-yr (IQR) | 51 (44.3-56.8) | 56 (53.3-59.8) | 57.5 (56.3-59) | 55 (53-60) | 54 (49-58) |
| Sex- male % (n) | 40 (4) | 60 (6) | 80 (8) | 70 (7) | 81 (13) |
| Antihypertensive medication use*- % (n) | 30 (3) | 40 (4) | 30 (3) | 30 (3) | 25 (4) |
| Diabetes- % (n) | 10 (1) | 0 | 0 | 30 (3) | 13 (2) |
| Statin use- % (n) | 20 (2) | 30 (3) | 20 (2) | 20 (2) | 13 (2) |
| Cause of death | - Cerebral bleeding: 40 (4)<br>- Accident: 30 (3) | - Cerebral bleeding: 70 (7)<br>- Cerebral occlusion: 10 (1) | - Cerebral bleeding: 20 (2)<br>- Myocardial infarction 30 (3) | - Cerebral bleeding: 18 (2)<br>- Myocardial infarction: | - Myocardial infarction: 88 (14)<br>- Cerebral ischemia: 6 (1) |
|  | -<br>Suffocation 10 (1)<br>- Cardiac<br>bleeding 10 (1)<br>-<br>Pulmonary embolus 10 (1) | -<br>Pneumonia : 10 (1)<br>-<br>Myocardial infarction 10 (1) | - Cardiac<br>arrest: 20 (2)<br>- Cerebral<br>occlusion: 30 (3) | 45 (5)<br>- Cardiac<br>arrest: 9 (1)<br>-<br>Pneumonia : 9 (1)<br>-<br>Ventricular<br>fibrillation: 9 (1)<br>-<br>Esophageal<br>varices 9 (1) | -<br>Cerebrovascular disease,<br>unspecified: 6 (1) |

\* Antihypertensive medications included a: including Angiotensin II receptor blockers, ACE-inhibitors, diuretics and calcium channel block.

### Pseudo-timeline of atherosclerotic lesion progression and plaque morphology

The successive stages of atherosclerotic lesion progression included in the timeline are defined on basis of clear changes in core morphology on the Movat staining (**Figure 1**).^5^ The earliest progressive lesion type, PIT, is characterized by the emergence of a largely acellular area of intimal extracellular lipid accumulations, the so-called ‘lipid pools’ (**Figure 1, A1/A2-PIT**).^5^ By definition, cholesterol clefts are absent in this lesion type. Emergence of (small) cholesterol clefts and necrotic core formation, consisting of cholesterol and cellular debris, hallmarks transition from PIT to EFA (**Figure 1, B1/B2-EFA**).^5^ Large cholesterol clefts, a more amorph necrotic core and a thick overlying cap (**Figure 1, C1/C2-LFA**) characterize LFA, the classic atherosclerotic lesion type.^5^ Thinning and ultimate disruption of the cap characterize the unstable lesion types: TCFA and PR. Classification of the latter two lesions is essentially based on respectively cap thickness (<65μm thick; **Figure 1, D1/D2-TCFA**), or disruption of the overlying cap with (partial) washout of the necrotic core and fibroin deposition (**Figure 1, E1/E2-PR**).^5^

**Figure 1:**
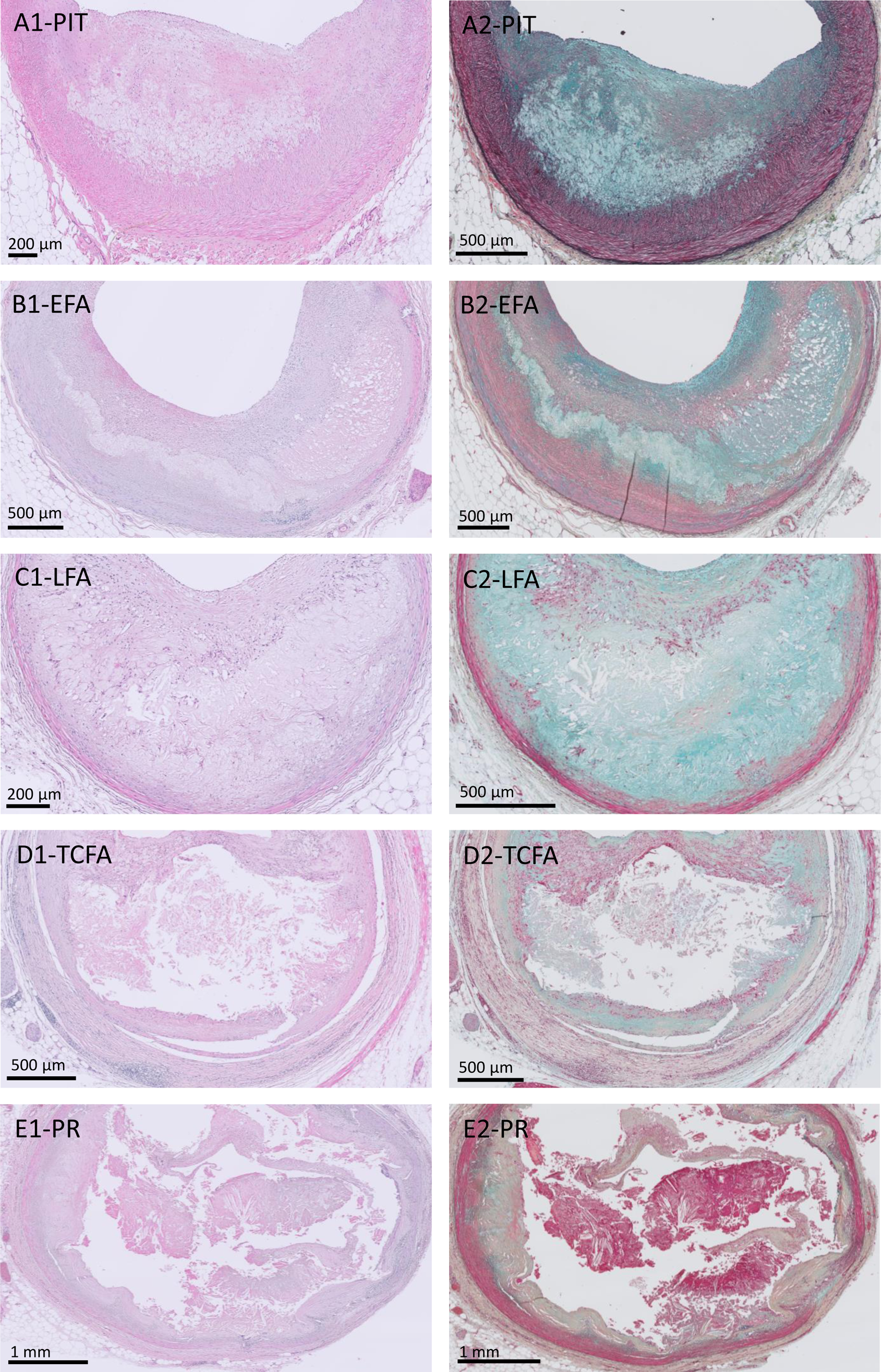
Histological summary of the changes in coronary plaque morphology during atherosclerotic progression (HE and Movat stainings of consecutive sections) Progressive atherosclerotic lesions are characterized by extracellular lipid accumulations, the so-called ‘lipid pools’,e.g. in pathological intimal thickening (PIT, A1-A2) that can converge into necrotic cores, e.g. in early fibroatheroma (EFA, B1-B2). Necrotic cores consist of extrcellular lipids, cholesterol crystals and necrotic debris. Progession into LFA (C1-C2) is characterized by expansion and organisation of the necrotic core. In thin cap fibroatheroa (TCFA, D1-D2), the necrotic core is usually even larger. In plaque ruptures (PR, E1-E2), the necrotic core components are partially washed away and the thrombus or remnants of the thrombus are in continuity with the underlying necrotic core. Color legend of Movat Pentachrome staining: *blue*, proteoglycans; *yellow*, collagen; *green/ochre*, indicate different proportions of proteoglycan/collagen colocalization; *black*, elastin; *red*, smooth muscle cells (SMC) or fibrinogen; *purple*, nuclei. Color legend of HE staining: *blue-purple*, cell nuclei; *pink*, extracellular matrix

### Changes in cap morphology during atherosclerosis progression

The pseudo-timeline shows clear changes in cap morphology during lesion progression (**Figure 2** and **Supplemental Figure 3**). A morphologically distinct cap is absent in PIT, and the lipid pools are covered by a thick, proteoglycan and mesenchymal cell rich intimal area, that is morphologically similar to that of the neighboring non-lesional intima. Atherosclerotic lesion progression is associated with a gradual transformation of the overlying intima into a fibrotic cap. The different aspects of the transformation process (matrix composition, cell/matrix ratio, thickness of the cap, collagen structure, mesenchymal cell orientation and nucleus morphology, leukocyte infiltration, neovascularisation, and cell death) are discussed in the following sections.

**Figure 2:**
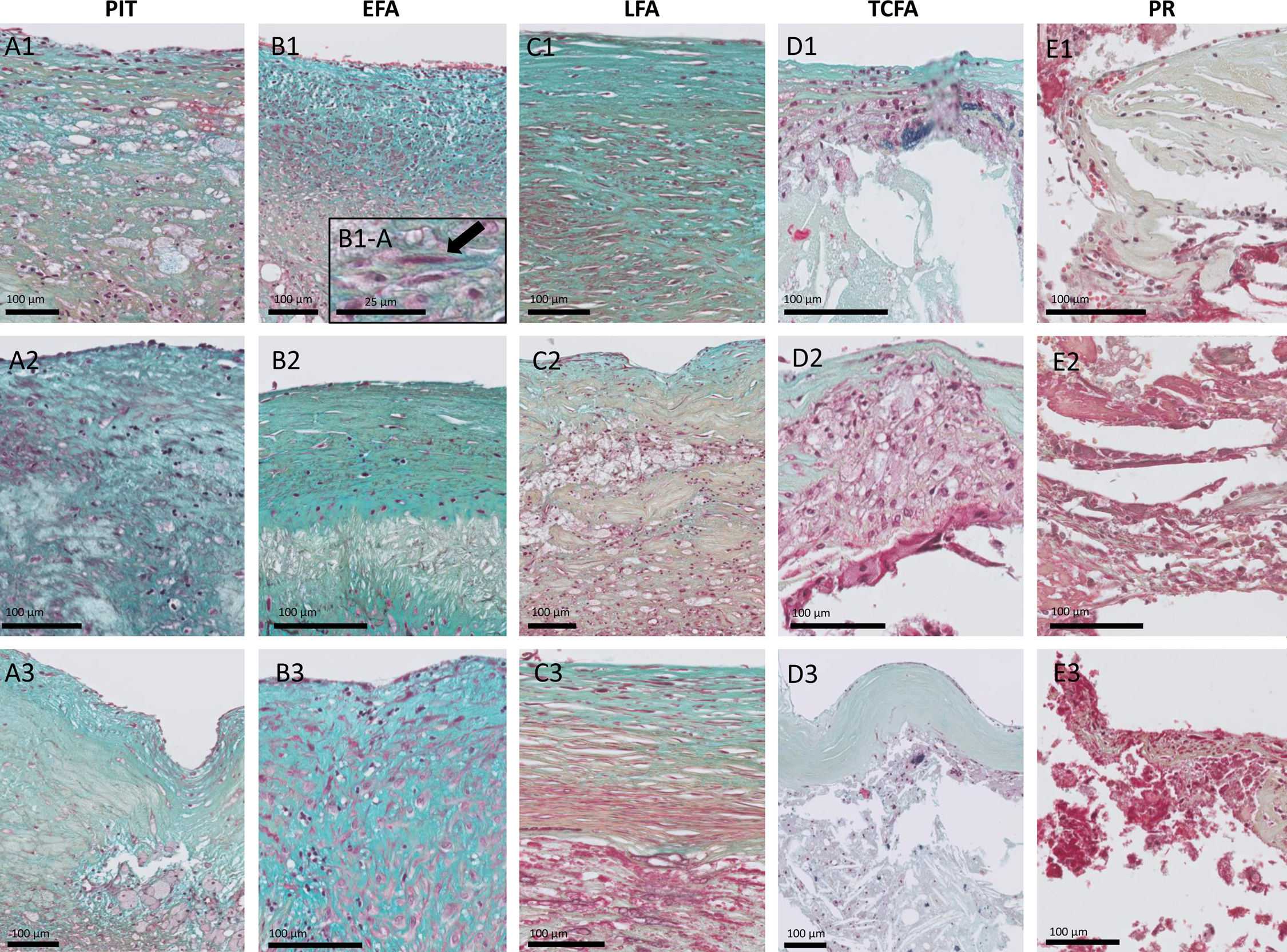
Histological overview (Movat Staining) of coronary cap morphology during progessive stages of atherosclerosis. The rows represent the different stages of progressive atherosclerotic lesions. The columns represent examples from three distinct patients illustrating the inter-patient variability. All images are at the same magnification, as such for lesions with a thicker cap, only part of the cap is represented. During early stages (PIT, A1-A3) of the atherosclerosis process, the thickened intima is mesenchymal cell rich (e.g. smooth muscle cells) and proteoglycan rich. Macrophages and foam cells may or may not diffusely infiltrate the thickened intima. A clear mesenchymal cell orientation is absent. Progession towards Early Fibroatheroma (EFA, B1-B3) is characterized by polarization of the matrix in the thickened intima (increased collagen content on the abluminal side covering the necrotic core (ochre hue) and a high proteoglycan/collagen ratio in the luminal aspect of the cap (blue hue)); a cap is emerging. Degrees of leukocyte content vary. Morphologic changes (nucleus elongation) are observed in a subset of the mesenchymal cells (B1-A). Compared to PIT, the mesenchymal cell orientation is more disorganized. Progression to a LFA (C1-C3) associates with a clear decrease in the cap’s mesenchymal cell density and a decrease in the proteoglycan/collagen ratio in both the luminal and abluminal aspect of the cap. Mesenchymal cells and nuclei are more elongated. Mesenchymal cells in the cap align with the collagen fibers, resulting in more organization. Leukocytes present mainly in the abluminal aspect of the cap. By definition, TCFA lesions (D1-D3) are characterized by a cap thickness of less than 65 µM. The ochre hue of the cap’s extracellular matrix in the Movat staining indicates a collagen dominated extracellular matrix. The mesenchymal cells are almost depleted. An exclusive finding for a subset of TCFA lesions is the emergence of ‘foam’ cells in the central portion of the cap (D1/D2). These cells mainly reside in the luminal portion of the cap rather than at the core aspect of the cap. Cap rupture (PR, E1-E3) is characterized by disruption of the thin, collagen dominated, severe mesenchymal cell depleted, cap. For the color legend of the Movat Pentachrome staining, please appreciate Figure 1. For a more detailed overview of cellular density in the cap, please appreciate Supplemental Figure 2.

### Pathological Intimal Thickening (PIT)

In PIT, the earliest progressive lesion type, the thickened vital intima overlying the lipid pools (the precursor of the cap, **Figure 2A, A1-A3**; and **Supplemental Figure 3, A1-A3**) resembles the adjacent thickened intima. The center section of the overlying intima was generally thicker than the lateral portions (**Supplemental Figure 4**). The predominant ‘longitudinal’ mesenchymal cell orientation that is characteristic of the ‘normal’ thickened intima was partly lost in the overlying section (**Figure 2A, A1-A3**).

The collagen fiber structure of the overlying intima in PIT lesions was variable, ranging from areas with a relative dense collagen fiber network to areas with breaks and thinned fibers (**Figure 3**).

**Figure 3:**
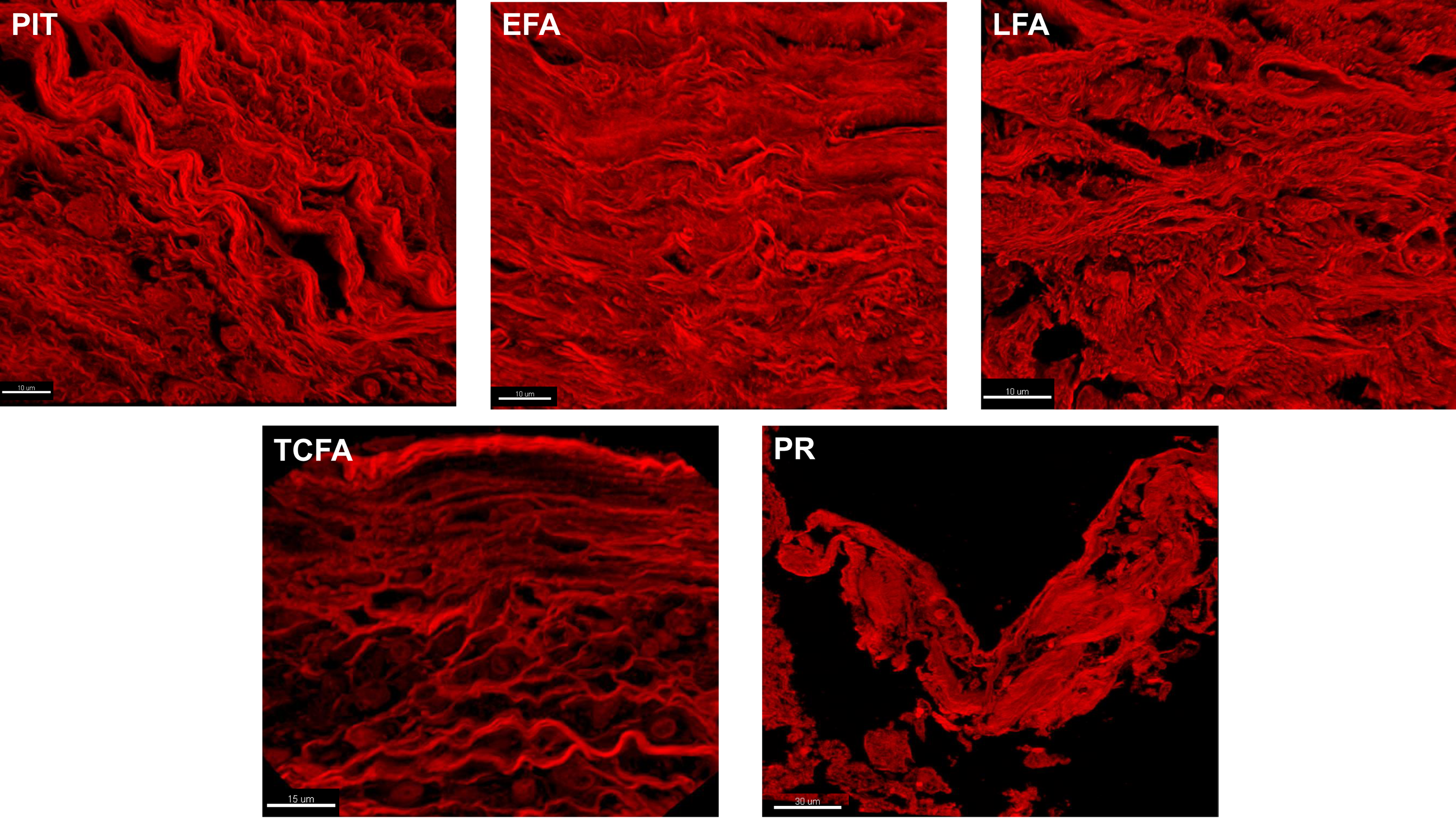
Collagen structure during atherogenic progression. 3D reconstructions (based on Z-stacks) of the collagen structures in the caps of progressive coronary atherogenic lesions (Progressive Intima Thickening to Plaque Rupture) were created based on confocal Z-stacks from 4μm sections. Thinning of and breaks in the collagen networks was observed during progression, leading to a brittle collagen structure in Plaque Ruptures.

While macrophages (CD68+/CD45+; **Supplemental Figure 5, A1**) primarily present in the areas adjacent to the lipid pools, abundant non-macrophage leukocytes (CD45+/CD68-; **Supplemental Figure 5, A2**) populate the intimal segment overlying the lipid pools. Very limited intimal neovascularization (microvessels) was observed in this early stage lesion (**Supplemental Figure 6A**). Less than 10% of the intimal leucocyte and of the mesenchymal cell population stained positively for the apoptosis marker Cleaved Caspase-3 (**Supplemental Figure 7A**), and for the pyroptosis marker Gasdermin-D (**Supplemental Figure 8A**).

### Early Fibroatheroma (EFA)

Transition to an EFA is accompanied by polarization of the intimal segment that overlyies the necrotic core into a distinct, cap-like structure (**Figure 2B, B1-B3**). Movat stainings indicated spatial diversity of the cap matrix: while a high proteoglycan/collagen ratio (reflected by the blueish color) was observed in the luminal aspect of the cap, the areas adjacent (‘abluminal side’) to the core were characterized by a decreased proteoglycan/collagen ratio (reflected by the ochre hue in the Movat staining). The cap/shoulder thickness ratios were variable (**Supplemental Figure 4**). The 3D collagen fiber reconstructions indicated more breaks compared to PIT (**Figure 3**), and clustering of collagen fibers, especially on the abluminal side. Changes in mesenchymal cell orientation were more pronounced compared to PIT; the majority of the mesenchymal cells had a circular orientation, rather than a longitudinal orientation. A subset of the mesenchymal cells showed clear morphologic changes with pronounced elongation of the cell and nucleus (**Figure 2B, B1-B3**; and **Supplemental Figure 3, B1-B3**).

Macrophages and foam cells (CD68+; **Supplemental Figure 5, B1**) mainly resided at the abluminal side of the cap, adjacent to, or confluencing with the necrotic core (**Supplemental Figure 5, B3**). Non-macrophage leukocytes (CD45+/CD68-, **Supplemental Figure 5, B2**) dominated the luminal sides of the cap.

Cap and shoulder neovascularization were both minimal; although microvessels were more abundant in the shoulder regions than in the cap (**Supplemental Figure 6, B1/B2**). The apoptosis index (%-cells positive for cleaved Caspase-3) and the pyroptosis index (%-cells positive for Gasdermin-D) of the mesenchymal cells in the cap was low (<10%; **Supplemental Figure 7B and Supplemental Figure 8B**). The apoptosis index of leukocytes in the cap was higher (10-50%, **Supplemental Figure 7B**); a phenomenon that was mainly due to the large proportion of foam cells being positive for the apoptosis marker cleaved Caspase-3. Less then 10% of the leukocytes in the cap stained positively for the pyroptosis marker Gasdermin-D (**Supplemental Figure 8B**).

### Late Fibroatheroma

Progression from an EFA to a LFA is accompanied by a clear changes in cap characteristics: i.e. clear decreases in the proteoglycan/collagen ratios in both the luminal and abluminal aspects of the cap (**Figure 2C, C1-C3**), emergence of coarse collagen bundles in the abluminal aspect of the cap (**Figure 3**), and a decrease in mesenchymal cell density (**Supplemental Figure 3, C1-C3**). Morphologically these observations are consistent with a transition towards a more fibrotic tissue phenotype.

Alignment of the mesenchymal cells with the collagen bundles resulted in a higher degree of organization than in the earlier lesion types. Progression to a LFA lesion type was also accompanied by further changes in mesenchymal cell morphology and orientation: cells and their nuclei became more elongated (**Figure 2C, C1-C3**; and **Supplemental Figure 3, C1-C3**).

A notable aspect for the LFA’s was the emergence fragile, elongated mesenchymal cells with delicate nuclei, and minimal cytoplasm that located in voids of the densely packed collagen (**Figure 4A**). These notable morphological changes bear close similarities with morphological changes previously reported for subsets of mesenchymal cells present in fibromas that were referred to as an “inanotic”10 cell type. Confocal imaging (**Figure 4B**) showed that these “inanotic” cells stained positively for mitochondria (SOD2+) and for Vimentin cytoskeletal filaments. In contrast, expression of the senescence marker p16INK4a was limited and in fact mainly confined to cells infiltrating the necrotic core, especially foam cells (**Supplemental Figure 9**).

**Figure 4:**
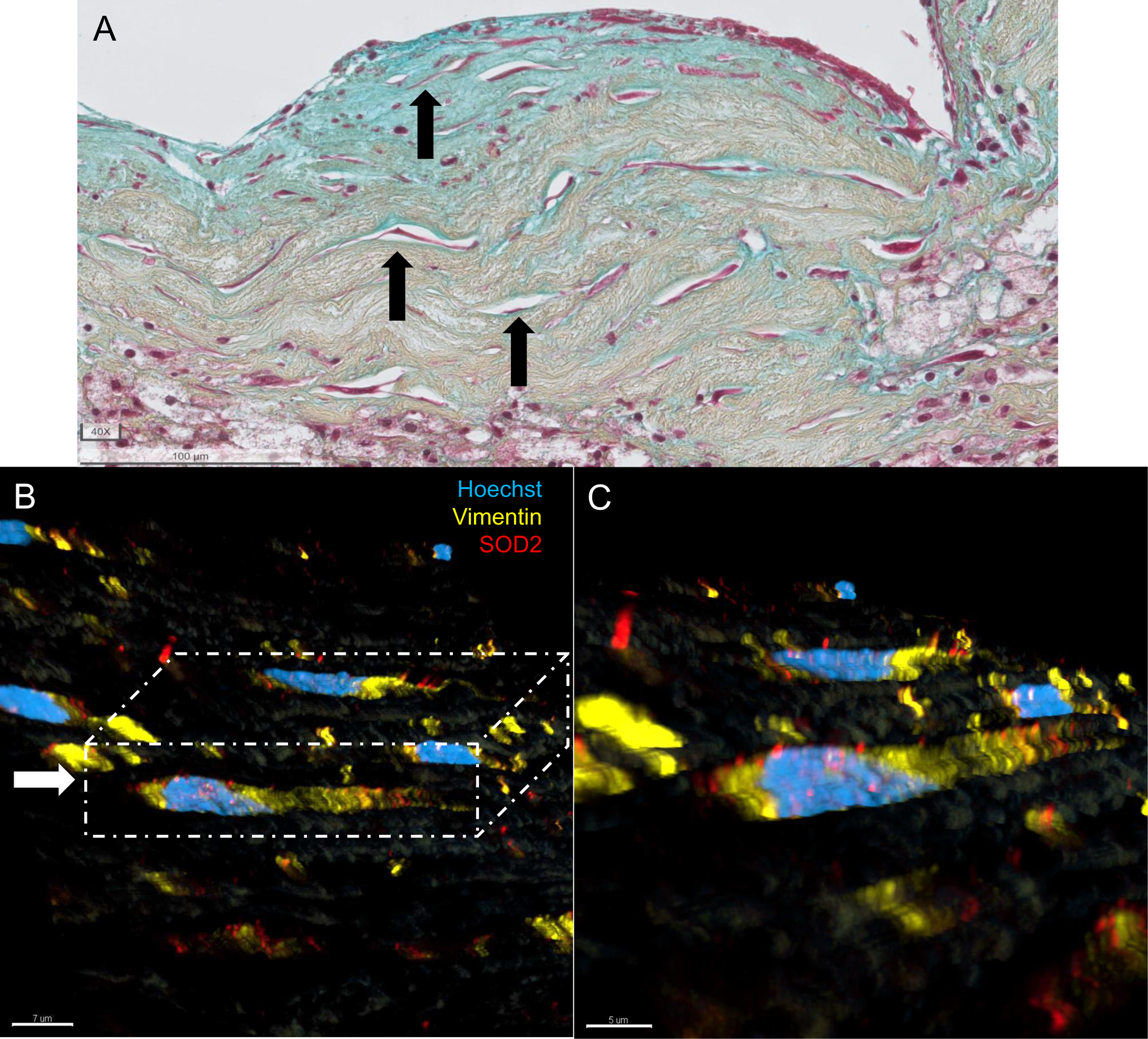
Mesenchymal Inanosis in advanced atherosclerotic lesions. Fragile cells with wispy elongated nuclei and minimal cytoplasm in densely packed fibrous areas (**A**) in advanced atherosclerotic lesions (here Late Fibroatheroma is shown) were observed. This appearance in fibromas has been described as inanosis. Confocal imaging delineated cellular composition (**B**): frail nuclei (Hoechst+, *blue*) were associated with mitochondria (SOD2+, *red*) and Vimentin filaments (key component of cytoskeleton, *yellow*). For the color legend of the Movat Pentachrome staining, please appreciate Figure 1.

Compared to the EFA lesions, the apoptosis index of mesenchymal cells in the cap was higher (10-50%, **Supplemental Figure 7C1**, and **Supplemental Figure 8C**) and in leukocytes comparable (10-50%; **Supplemental Figure 7C2).** The pyroptosis indices for both leukocytes and mesenchymal cells remained low (<10% respectively **Supplemental Figure 7C and 8C**).

A clear gradient in cap leukocytic infiltration (**Supplemental Figure 5, C1**) was observed with both macrophages and non-macrophage leukocytes mainly residing in the abluminal aspect of the cap (**Supplemental Figure 5, C2/C3**). Cap neovascularization remained minimal, and neovascularization was essentially confined to the shoulder regions (**Supplemental Figure 6-C1**).

### Thin Cap Fibroatheroma (TFCA)

By definition, TCFA lesions are defined in the revised AHA classification by a cap thickness less than 65 µM (**Supplemental Figure 4**). Qualitatively, these thin caps are characterized by a coarse collagenous matrix (ochre hue in the Movat staining, **Figure 2D, D1-D3**). Compared to LFA, the collagen fibers appeared thinner, and voids were common (**Figure 3**, arrows in TCFA). The thin cap was almost devoid of mesenchymal cells (**Supplemental Figure 3, D1-D3**), and the apoptosis and pyroptosis indices of the remaining mesenchymal cells were high (>80%, **Supplemental Figure 7D** and **Supplemental Figure 8D**).

A notable observation in two thirds of the TCFA lesions was the infiltration of foam cells in the central portion of the cap (**Figure 2D, D1-D3**). These cells resided both in the luminal portion of the cap and at the core aspect of the cap. Further characterization (**Supplemental Figure 5, D1-D2**) of these foam cells suggested heterogeneous identities with both macrophage origin (CD45+/CD68+/Vim-; **Supplemental Figure 5, D3-D7**) as well as mesenchymal (CD45-/CD68+/Vim+; **Supplemental Figure 5, D3-D7**) origin. The majority of these foam cells (>80%) stained positively for the pyroptosis marker GSDM-D (**Supplemental Figure 8D**); and for the apoptosis marker Active Caspase-3 (50-80%; **Supplemental Figure 7D**). Lesion neovascularization remained stable, and limited to the shoulder regions. (**Supplemental Figure 6, D1**).

### Plaque Rupture (PR)

Cap rupture was accompanied by disruption of the cap, and (partial) discharge of core contents leading to a (partial) collapse of the cap and core (**Supplemental Figure 10**). Varying degrees of fibrin deposition can be present (**Figure 2E, E1-E3; Figure 3**). Rupture mainly occurred in the lateral sections of the cap (in 7 of 14 cases), but was also observed for the central (4 of 14 cases) or shoulder (3 of 14 cases) sections of the cap (**Supplemental Figure 11**).

## Discussion

A necrotic core with an overlaying fibrotic cap is the archetypical atherosclerotic lesion.^11^ Rupture (destabilization) of this cap is almost universally considered the key trigger of the acute, thrombotic complications of atherosclerotic disease.^12^ Although detailed classification schemes of morphological changes in atherosclerotic plaque development exist^5^, less attention has been paid to qualitative changes of the atherosclerotic cap during progression. The pseudo-time line of coronary atherosclerotic lesion development, maturation and destabilization established in this study, shows an almost uniform pattern of cap maturation and destabilization, with progressive fibrotic changes leading up to terminal failure (rupture).

The concept of plaque rupture as the primary trigger of acute thrombotic events has found almost universal acceptance, and its putative underlying mechanisms have been summarized in numerous reviews. The current main concept is that cap destabilization is driven by an imbalance between excess protease activity (released from macrophages and mesenchymal cells), and impaired ECM production as result of mesenchymal cell death; with a chronic, localized inflammatory response as the overarching theme. ^13,14,15^ Further aspects implemented include mesenchymal cell senescence^16^, impaired efferocytosis (clearance of apoptotic cells by phagocytes) leading to accumulation of necrotic cells^17^, defective resolution of inflammation^18^, and plaque neoangiogenesis^19^. The latter is thought to sustain leucocyte influx leukocytes, and to promote plaque progression through leakage of erythrocytes and thereby accumulation of cholesterol and erythrocyte-derived phospholipids.^19^

The above concepts are essentially based on extrapolation of plaque characteristics of individuals who died from myocardial infarction, or plaque material removed during a carotid endarterectomy, as well as on observations from experimental models of plaque rupture. However, because the clinical samples essentially reflect the post-rupture condition, they are mainly representative for the final stages of the destabilization process, and may not adequately discriminate between cause or consequence. On the same token, interpretation of the experimental rupture models is hampered by the lack of validation, and it remains unclear to what extent they adequately represent aspects of clinical cap maturation and destabilization.^8^

Relying on a repository of coronary artery specimen from young to middle aged tissue donors (maximum age 70 years), and using the revised AHA (Virmani) classification^5^ for guidance, we were able to construct a pseudo-timeline of the atherosclerotic process in order to study the qualitative dynamics of cap formation and destabilization. This timeline was dictated by the changes in core morphology and quantitative changes in cap thickness. The primary focus was on matrix and mesenchymal cell dynamics. The cellular aspects of the innate and adaptive immune responses during the human atherosclerotic process have been extensively evaluated and reported in earlier series of studies.^20,21,22^

This evaluation shows that cap formation is intrinsically linked to the progression of atherosclerosis. A distinct cap structure is missing in the PIT, the earliest progressive atherosclerotic lesion, in fact the tissue overlaying the lipid pool was morphologically very similar to the adjacent thickened intima. Transition to EFA is accompanied by polarization of the intima overlying the developing necrotic core, creating a discrete cap. Progression to a LFA is accompanied by progressive fibrotic changes in the cap characteristics with a clear decline in the proteoglycan/collagen ratio and mesenchymal cell density. Although fibrotic changes are often interpreted as a sign of plaque stability^23,24^, the question arises whether this assumption is justified. In fact, the data in this study suggest that the fibrotic changes are a prelude for destabilization. Reconstructions of the collagen fibers indicate clear changes in matrix structure during atherogenic progression. This is a paradoxical aspect in fibrosis: although the deposition of a dense collagenous matrix may suggest a more stable matrix^25^, the loss of network structure in fibrosis actually results in a more brittle matrix.^26^ One could speculate that the collagen voids seen in the cap of LFA and TCFA lesions are in fact an artifact and reflect dissociation of the brittle collagen bundles caused by shrinking and expansion of the tissue during tissue procurement (the dehydration-rehydration steps in the tissue procurement for paraffin embedding and later in staining processes).^27^

Transition to a more hostile micro-environment is supported by an increase of cell death during atherogenic progression, and the emergence of cells with morphological similarities similar to those described as inanotic cells in the context of fibroid myomas.^10^ Inanosis is a type of cell death resulting from gradual nutritional deprivation (inanition), without a phagocytic reaction, but rather dissolution of the cell (reclamation).^10^ Cells with similar morphological characteristics are abundantly present in advanced atherosclerotic caps.

Based on the processes described for the myoma’s, we propose that a similar micro-milieu may occur in the central portion of the atherosclerotic cap. The stenosis accompanying the atherosclerotic lesions will result in an accelerated and turbulent luminal blood flow, further interfering with the normally already minimal gas and nutrients exchange between the coronary blood flow and the adjacent tissue, whereas the underlying necrotic core precludes supply from below. As a result, the cap will largely rely on transport from the shoulder regions. As a consequence, an enlarging core during lesion progression results in progressive larger distances between the central portion of the cap and its supporting structures (e.g. the vasa vasorum in the media and adventitia; and in the shoulder regions).

Conclusions for plaque neovascularization were in line with previous studies^28^, and show that neovascularization is mainly confined to the shoulder areas from early stages on, and that neovascularization of the central portions of the cap is minimal. Polarization of the luminal and abluminal aspects from the cap point to possible direct toxic interaction between the necrotic core contents and the adjacent tissue.^29^ Progressive cap thinning will expose larger proportions of the cap to this interaction.

The deprived and hostile micro-environment may ultimately lead to exhaustion of the mesenchymal cell population and thus a highly fibrotic cap structure that is unable to keep up with the progressively growing lesion. This aspect appears rather underappreciated in the existing literature, and could explain the disappointing results from (pre)clinical studies^30^ aimed at cap stabilization.

Whilst foam cells were virtually absent in the caps of EFA’s and LFA’s, foam cells populate the central portion cap in the majority of the TCFA lesions. One could speculate that this phenomenon relates to a change in matrix structure towards a more promigratory phenotype. Alternatively, and non-exclusively, the emergence of foam cells may be related to chemotactic signals that appear to precede plaque rupture. In a previous study we observed aspecific chemokine signature characterized by CXCL-13 expression and CXCL-13-mediated responses that appeared exclusive for TCFA’s.^21^ It could be speculated that the foam cell infiltration involves a specific chemokine signal. The observed foam cell infiltration might be involved in the terminal destabilization process, either directly by matrix displacement, or indirectly by the release of plasminogen activators, cathepsins, and matrix metalloproteinases, that are able to degrade collagenous matrix.^31^

Rupture was predominantly observed in the lateral sections in the cap, and not as generally reported in the shoulder regions^32,33^, or in the mid portion of the fibrous cap (as reported for plaque rupture during exercise).^5^ This apparent contrast could be an artifact that was caused by the rather ambiguous definition of the cap and shoulder areas.^34,35^ Alternatively, cap rupture and washout of the core contents will result in a pressure drop, and recoil of the cap and inversion of the shoulder regions, resulting in a distorted picture of the preceding situation.

Although this study depicts a consistent picture, it has several limitations. Firstly, since this is an observational study, no conclusions about causality or pathogenesis (e.g. foam cell infiltration in TCFA’s) could be drawn. Furthermore, the study population was dictated by the inclusion criteria for donation. As a result, the population was relatively young, non-diabetic and without chronic kidney disease. This aspect may explain the minimal incidence of plaque erosions^36^ observed in our tissue repository. Hence, observations from this study may not apply for plaque erosions.

The upper age limits for donation may contribute to the low proportion of women with an advanced stage atherosclerosis (LFA, TCFA and PR) in this study, impeding analysis of sex-specific characteristics. In fact, the PROSPECT trial^37^ showed that women are generally older when they develop an acute coronary syndrome. The trial also showed that women more often develop an acute coronary syndrome due to plaque erosion rather than plaque rupture; an aspect that may further contribute to the low proportion of women with an advanced lesion in this study.

Another possible contributing aspect to the low prevalence of advanced lesions in women, that cannot be explored in our observational study, is that the lesion destabilization process itself in women could be faster than in men. This aspect may introduce an artifact in cross-sectional studies.

In conclusion, this study characterizes the development of an unstable cap as (a possibly inevitable) consequence of a degenerative process leading to a fibrotic matrix and exhaustion of the mesenchymal cell population. It provides a unique framework for further studies aimed at reversing the degenerative process (e.g. fibrosis) and whether the destabilization-associated chemokine signature is causatively involved in the destabilization process.

## Nonstandard Abbreviations and Acronyms

EFA: Early Fibroatheroma
LFA: Late Fibroatheroma
LDL: Low-density lipoprotein
PIT: Pathological Intimal Thickening
PR: Plaque Rupture
TCFA: Thin Cap Fibroatheroma

## Acknowledgments

None.

